# High-throughput proteome profiling with low variation in a multi-center study using dia-PASEF

**DOI:** 10.1101/2024.05.29.596405

**Authors:** Stephanie Kaspar-Schoenefeld, Jonathan R. Krieger, Claudia Martelli, Ann-Christine König, Stefanie Hauck, Sebastian Johansson, Axel Karger, Uli Ohmayer, Matteo Pecoraro, Stefan Tenzer, Ute Distler, Sophie Braga-Lagache, Phillip Strohmidel, Lisa Abel, Raphael Schuster, Georg Kliewer, Tobias Kroniger, Laura Heikaus, Diego Assis, Torsten Mueller, Daniel Hornburg

## Abstract

High throughput proteomics is gaining increasing traction as it facilitates screening of large sample cohorts required in clinical research and systems biology studies. Recent developments in mass spectrometry-based proteomics resulted in improved hardware and software providing deep proteome coverage, robustness, and scale accessible to a wide range of laboratories. Here, we benchmark dia-PASEF, a data-independent acquisition scheme that integrates trapped ion mobility with high scan speed, with a high-resolution time-of-flight mass analyzer (timsTOF HT) for the deep proteome analysis of a human cell line applying short 5-minute gradients. To show intra-and interlaboratory reproducibility, we performed a multi-laboratory study including 11 sites. We demonstrate that on average 7,072 protein groups and 99,835 peptides were identified in human chronic myelogenous leukemia cells on the timsTOF HT with low variation. Our results underline that dia-PASEF data acquisition combined with reproducible chromatography enables high robustness and data consistency across instruments and laboratories, which is a prerequisite for translational biomedical insights.

## Introduction

Mass spectrometry (MS)-based proteomics holds tremendous potential for gaining translational biomedical insights inaccessible with next generation sequencing (NGS). Despite this advantage, proteomics has historically lagged behind NGS in scale and depth owing to the higher complexity of proteomics workflows required to capture the complex biochemistry of proteins and peptides. In recent years, MS-based proteomics has experienced fundamental advancements in instrumentation, acquisition, and software, with the promise to match and ultimately exceed NGS data generation.

The introduction of Trapped Ion Mobility Spectrometry (TIMS, [2]) technology is pivotal for greater proteomics depth as it not only enables highly efficient sampling of molecules in a sample (Parallel Accumulation Serial Fragmentation (PASEF, [3,4])) but also provides an additional dimension of separation as well as molecular characterization (collisional cross-section). This additional dimension reduces complexity by enabling separation of peptides that would be otherwise co-fragmented. TIMS thereby provides higher confidence and cleaner MS and MS/MS-spectra, and further increases sensitivity due to the space and time focusing of ions eluting from the ion mobility device. Using Parallel Accumulation Serial Fragmentation (PASEF, [3,4]) up to 30 precursors are addressable in a 100 ms TIMS scan, resulting in a sequencing up to 300 Hz. To gain the maximum degree of reproducibility data independent acquisition (DIA) approaches have attracted attention as stochastic changes resulting from abundance driven precursor selection as it is the case for data dependent acquisition (DDA) schemes have been eliminated. Instead, in DIA all ions in a certain mass range are co-isolated by the quadrupole and hence co-fragmented [5]. The resulting high spectral complexity is a major challenge of DIA, that is partly overcome by the introduction of TIMS to the concept of DIA and the development of dia-PASEF. The additional ion mobility separation reduces spectral complexity and the synchronization of TIMS with the quadrupole improves the ion sampling efficiency, compared to conventional DIA [1]. dia-PASEF has been shown to be very powerful regarding proteome coverage [1], in ultra-low sample amount studies like single cell proteomics [6] and immunopeptidomics [7] as well as in the study of post-translational modifications [8].

Take together, dia-PASEF is particularly well-suited for large cohort studies, which require robust and highly reproducible workflows in conjunction with short runtimes and high proteome coverage. Since population scale data may be integrated across multiple years and study centers, these performance metrics must be validated across laboratories and instruments.

Here we describe a diaPASEF acquisition scheme for a very short gradient length (5-minute active gradient) enabling the acquisition of ∼140 samples per day. The protein and peptide identification rates and intra-and interlaboratory reproducibility were evaluated across eleven independent laboratories. Our investigation not only highlights the potential of the timsTOF platform but also underlines its pivotal role in propelling proteomics research towards accessing more and larger sample cohorts.

## Material & Methods

### Sample preparation

Tryptic digest of a human cell lysate (K562, Promega) was dissolved in 500 µl 0.1% formic acid (FA) to a final concentration of 200 ng/µl. In-house HeLa and HEK digest were prepared as previously described ([9],[10]).

### LC-MS acquisition

LC-MS/MS analysis was performed using the same setup in all participating laboratories: a timsTOF HT mass spectrometer was coupled to a nanoElute 2 nanoLC system via a CaptiveSpray ionization source (all from Bruker Daltonics). One µl of prepared sample (corresponding to 200 ng was loaded directly on a 5 cm C18 column (75 µm inner diameter, 1.7 µm particle size, Aurora Rapid, IonOpticks). Peptides were separated using a 5 min gradient (from 3% to 26% buffer B in 4 min and from 26% to 40% in 1 min, buffer A: 0.1% formic acid, buffer B: 0.1% formic acid in acetonitrile) at a flow rate of 500 nl/min and a column temperature of 50°C. For washing the column, the organic solvent was increased to 90% buffer B in 0.2 min and maintained for 1.8 min.

For the dia-PASEF acquisition, a window placement scheme optimized via the py-diAID tool (https://pypi.org/project/pydiaid, [11]) was used. The resulting dia-PASEF method had an average window size of 25.5 Da (minimum: 12.27 Da, maximum: 122.81 Da) and consisted of 12 frames with 3 mass windows per frame (50 ms) resulting in a coverage of the m/z range from 350 to 1250 Da and a cycle time of 0.7 seconds (including one MS1 frame, Fig. 1, Supplementary Data 1). The TIMS ion mobility range was set from 1/K_0_ = 1.35 to 0.7 Vs cm^2^. Ten replicates were acquired in 11 different laboratories.

**Figure 1:**
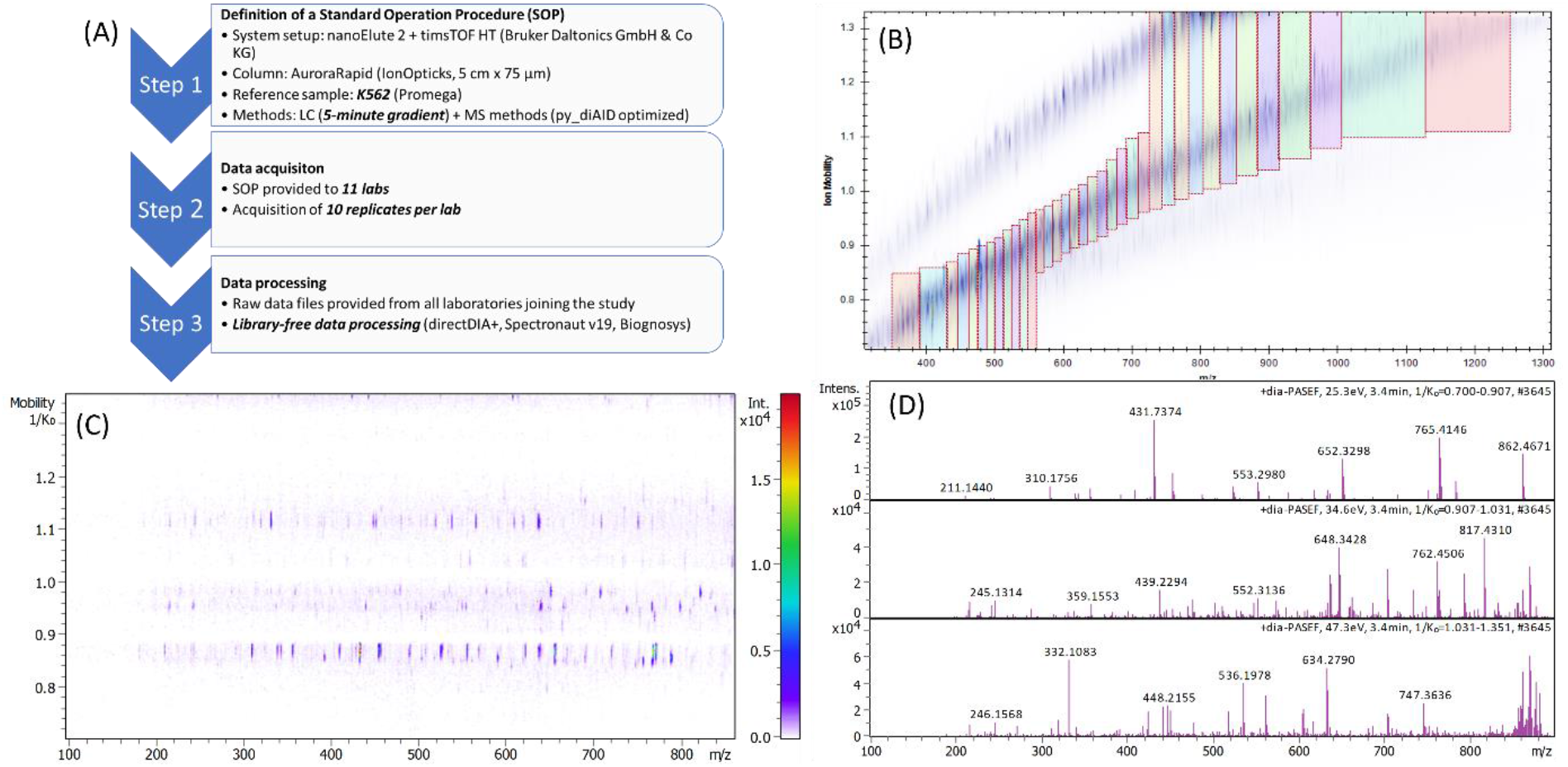
High-throughput proteome profiling in a multi-center study. (A) Detailed study overview (B) Optimized dia-PASEF made use of 3 mass windows per frame. An example frame is shown in (C) together with the corresponding MS/MS spectra (D).

### Data processing

dia-PASEF data files were analyzed in Spectronaut (v19, Biognosys) using library-free data processing (directDIA+). Human Uniprot fasta file (downloaded from https://www.uniprot.org/uniprotkb on 2023/03/18) was used for direct database identification from the dia-PASEF runs (with oxidation considered as a variable modification and cysteine modification of +57 Da as a fixed modification). False discovery rate (FDR) was controlled at 1% for peptide and protein level. All other parameters were kept at Spectronaut defaults settings.

## Results

### Optimized dia-PASEF method for high-throughput proteomics analysis of complex samples using a 5-minute gradient

Technical improvements have made MS-based proteomics amenable to robust, deep and high-throughput biological studies. We evaluated the proteomics performance on a timsTOF HT for a short 5-minute active gradient. When running such short gradients, the applied DIA methods’ cycle time must match the fast chromatography (narrow peaks of eluting peptides) to sample sufficient data points across the peaks to enable good quantification. Using dia-PASEF on the timsTOF HT platform allows optimal window placing to ensure fragmentation of all theoretical precursors, while maintaining coverage of the chromatographic peak. Here we used py_diAID [11], which automatically adjusts the isolation window width to the precursor density and optimizes the isolation window positioning and design in the mass to ion mobility space. By encompassing three mass windows per PASEF frame and implementing brief accumulation and ramp times of 50 ms on the TIMS device, we attained a cycle time of 0.73 s, including 1 MS1 scan. The chromatographic peaks had a median width of about 3 s. Thus, the applied dia-PASEF method resulted in an average peak coverage of 4 to 5 data points in a 5-minute gradient.

### Multi-laboratory study reveals high reproducibility at scale

For clinical proteomics, robust and fast analysis with excellent interlaboratory consistency of several hundreds of samples per day is highly desirable and mandatory for some study goals such as population scale and longitudinal profiling. It has been previously shown that dia-PASEF provides reproducible peptide and protein quantification with low coefficient of variation (CV) values for peptide and protein abundances across multiple injections [1]. Here, we investigated the reproducibility of the presented workflow across instruments in different laboratories. A commercially available reference sample (K562, Promega) was provided to 11 laboratories geographically located throughout Europe and North America. All laboratories followed the same protocol regarding sample preparation, instrument, and method setup (Figure 1A) to evaluate the number and reproducibility of peptide and protein group identifications between the participating labs. Ten replicates were acquired per site resulting in a data set consisting of 110 files. From the 5-minute gradient, on average 7,072 protein groups and 99,835 peptides were identified at 1% FDR using library-free data processing (Figure 2A and B). In total, 7,228 protein groups and 124,064 peptides were identified across the entire data set. This demonstrates the high depth of coverage and excellent protein identification consistency of the timsTOF platform for high throughput applications and large-scale cross laboratory comparable proteomics studies.

**Figure 2:**
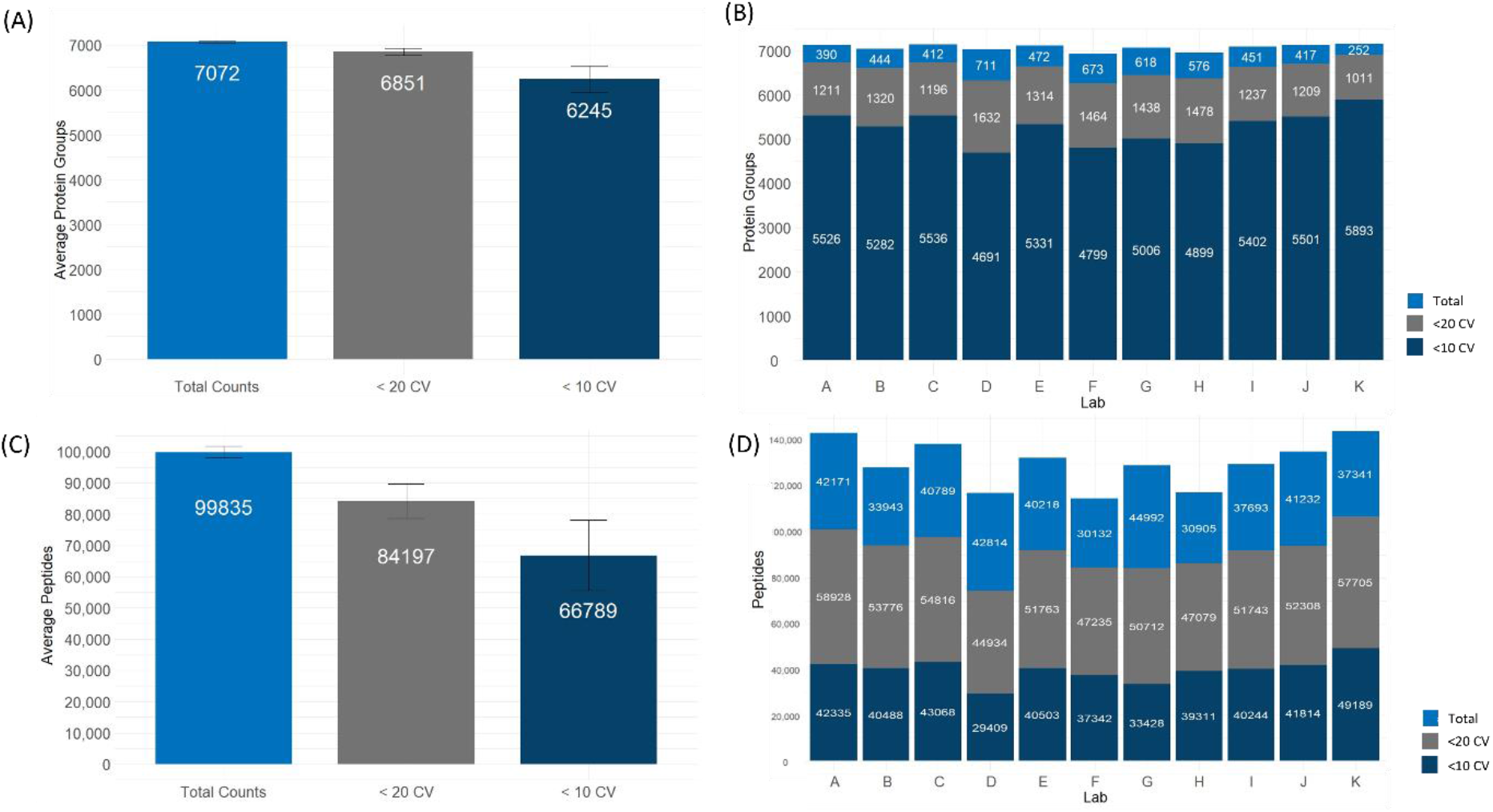
Number of identified protein groups from analysis of 200 ng K562 using a 5-minute gradient on the timsTOF HT with a py_diAID-optimized dia-PASEF method. (A) Average number of identified protein groups across all 110 data sets (11 labs, 10 replicates per lab) and number of quantified protein groups with CV values below 20% (grey) and 10% (dark blue). (B) Number of identified protein groups per lab (total, quantified with CV values below 20% and 10%). (C) Average number of peptides across all 110 data sets (11 labs, 10 replicates per lab) and number of quantified peptides with CV values below 20% (grey) and 10% (dark blue). (D) Number of identified peptides per lab (total, quantified with CV values below 20 and 10%).

Achieving a high number of proteins and peptides within a short measurement time is important, but identifications without sufficient quantification will have little value for addressing most scientific questions. Detecting molecular association with phenotypes, effect size, number of samples, and precision of quantification are determining factors.

The sequential elution of condensed ion packages from the TIMS device allows for even more efficient ion usage by increasing the effective duty cycle [1] and signal quality. The number of proteins identified with CV values below 20% and 10% ranged from 96% to 88%, respectively, across all 110 data sets. Average CV values for each lab on protein group level are well below 10% for the 10 replicates (Figure 3). The median CV value across the complete experiment (including all 110 runs) was at 12.1% on protein group level.

**Figure 3:**
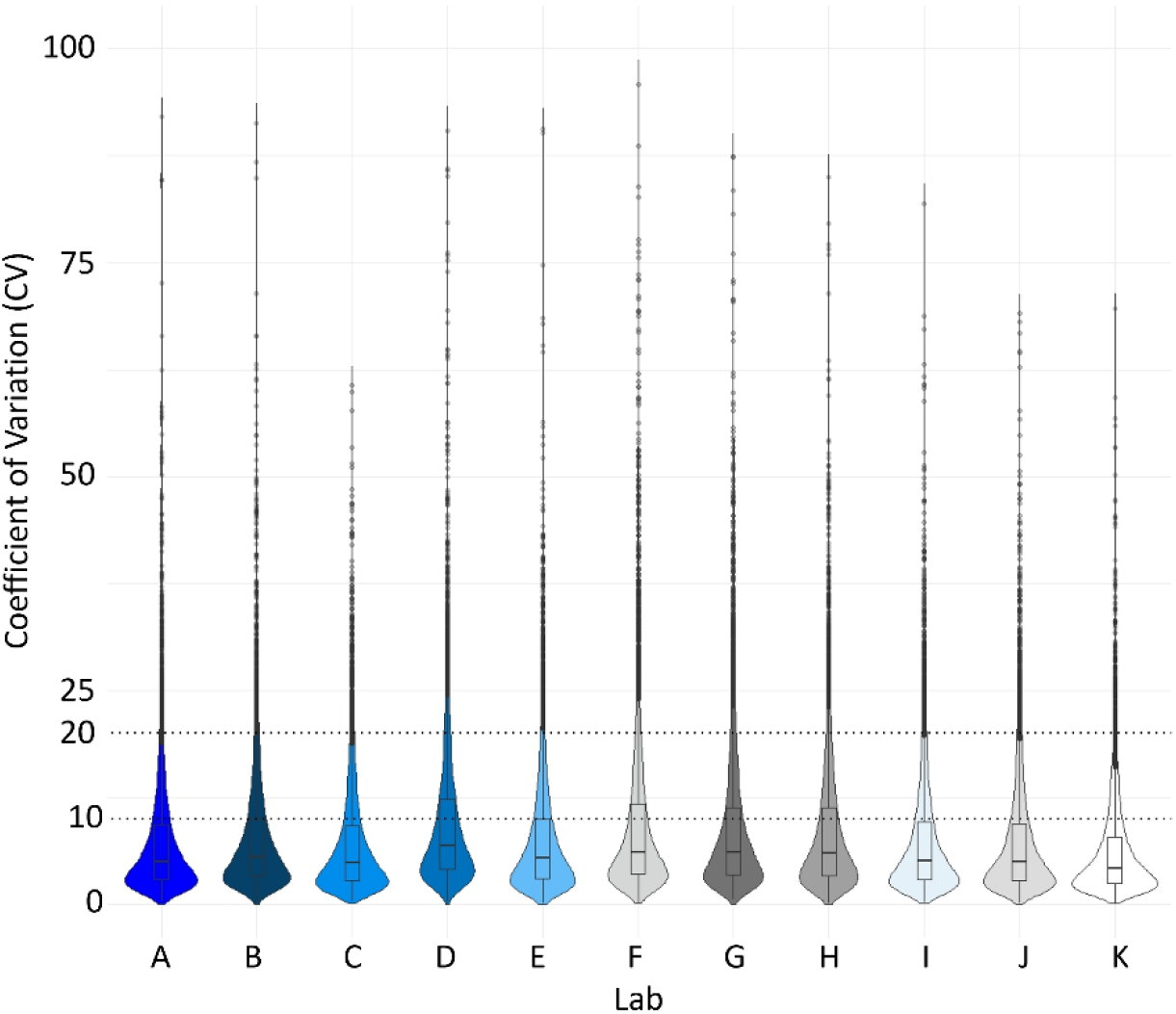
Coefficient of variation between the 10 replicates per laboratory on protein group level as calculated by the data processing software.

In total, 4616 protein groups have been identified in all 10 replicates of all labs participating in this study (corresponding to 110 data sets).

### Accessing the performance on different cell lines

The outcome of proteomics experiments is strongly dependent on the sample type used. In our study, a commercially available standard has been analyzed to compare the performance of dia-PASEF across different labs for high throughput proteomics. In addition to those measurements, we performed a comparison of K562 (Promega), HeLa, and HEK (in-house digests, respectively) tryptic digests using the setup as described above. On average 6,618 protein groups were identified from ten replicate injections of K562, whereas 7,444 and 6,908 protein groups have been identified for HEK and HeLa digests, respectively (Figure 4A). Interestingly, HeLa provided the lowest number of identified peptides (81,637), and HEK cell lines resulted in the identification of 30% more peptides (108,119, Figure 4B).

**Figure 4:**
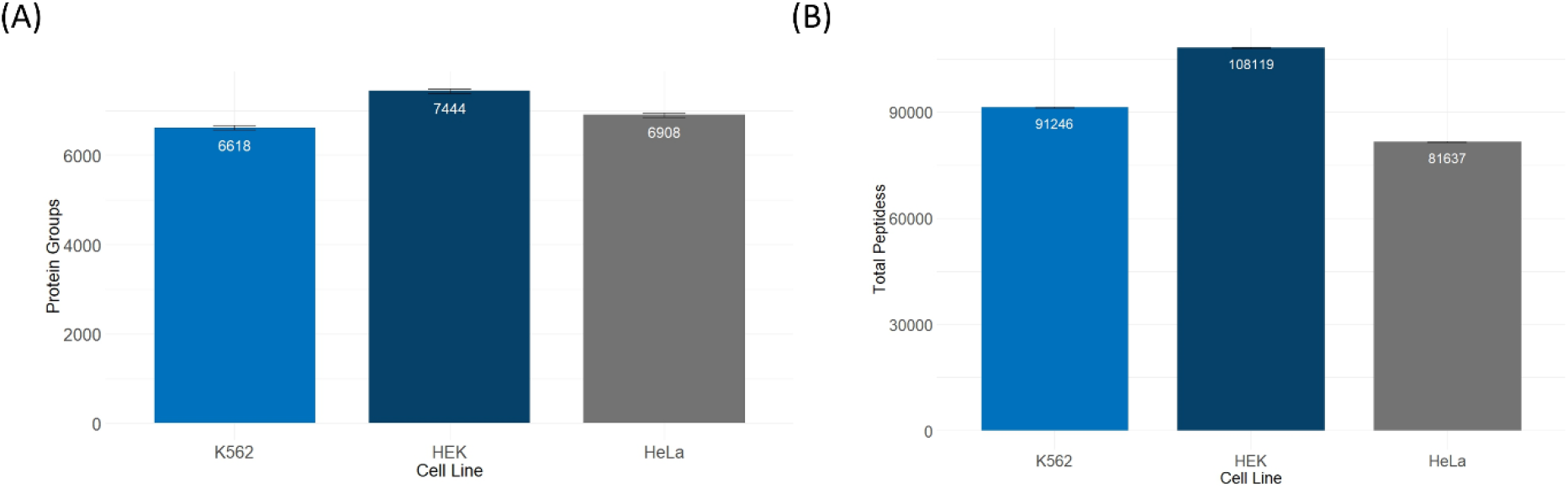
Average number of (A) identified protein groups and (B) peptides from different cell lines analyzed using a 5-minute gradient.

## Discussion

For the application of proteomics approaches in clinical and basic research, robustness and reproducibility are important features together with data completeness [13]. Whereas it is well-known that DIA delivers highly reproducible results not being biased by a precursor selection algorithm, only few studies focus on the feasibility to generate reproducible results across different labs for high throughput applications [14]. Improving throughput while maintaining high coverage and quantitative precision facilitates proteomics applications for large-scale biological experiments. In this study, we investigated the inter-and intra-laboratory reproducibility and overall performance of a dia-PASEF acquisition scheme for proteome analysis using noticeably short gradients of 5 minutes.

Laboratories participating in this study all used the same setup including reference sample, sample preparation, column, methods, and LC-MS system. Deploying an optimized dia-PASEF method resulted in excellent proteome coverage while maintaining good quantitation due to achieved peak coverage. The data from 11 different laboratories revealed an excellent average number of identified protein groups of 7072 and 99,853 peptides when measuring a commercially available reverence sample (K562). These results underline the utility of dia-PASEF for high-throughput deep proteomics providing state-of-the art results [15], enabling fast and comprehensive proteome profiling in biological studies. In total, 4616 protein groups were detected across all sites, indicating a high level of data completeness even across the 110 runs (11 laboratories with 10 replicates each).

A key advantage of dia-PASEF acquisition on the timsTOF platform are low peptide and protein group CV values between technical replicates across all 110 data sets underlining the high consistency of results obtainable using dia-PASEF on the timsTOF HT system. Therefore, the presented approach reveals highly reproducible results being an important prerequisite for large scale studies.

In this study, we could also show the dependency of the number of identified protein groups and precursors on the human cell type used. In our study, HEK provided the highest identification rates with 7,444 protein groups and 108,119 peptides on average from 200 ng sample injections. These findings underline the importance of using the same sample types when LC-MS workflows across different laboratories.

## Conclusion

High throughput proteomics requires a highly robust LC-MS system capable of handling the large volume of samples processed while maintaining stable performance over time without manual interference.

Here we demonstrate that dia-PASEF on timsTOF HT enables high-throughput, high-performance measurements of proteomics samples in a highly reproducible manner even with short gradients. More than 7,000 protein groups from approximately 100,000 peptides can be identified on average in a 5-minute gradient across 11 different labs.dia-PASEF results in excellent and reproducible multi-laboratory performance making the presented workflow ideally suitable for routine proteomics applications.

## Supporting information

S1_SOP_High-throughput proteome profiling with low variation in a multi-center study using dia-PASEF

S2_diaParametersfile

S3_ProteinGroupoutputtable

## Supplemental File Information

**File S1**: Standard Operating Procedures as provided to the participating laboratories.

**File S2**: diaParameters.txt file of the applied dia-PASEF method.

**File S3**: Protein Group output table.

