## Supplementary material for "High-throughput proteome profiling with low variation in a multi-center study using dia-PASEF": S1_SOP_High-throughput proteome profiling with low variation in a multi-center study using dia-PASEF

**Standard Operating Procedures for "High-throughput proteome profiling with low variation in a multi-center study using dia-PASEF"**

1. Dissolve the cell lysate (K562, V6951, Promega) with 500 µL 0.1% formic acid and transfer it into an autosampler vial

2. Make sure all solvents (A: 0.1% formic acid in water, B: 0.1% formic acid in Acetonitrile) and transfer liquid in the nanoElute2 system are fresh and run the "preparation" procedure

3. Connect the provided column (Aurora Rapid75 CSI 5cm, 5cmx75um, AUR3-5075C18-CSI, IonOpticks)

4. Run the "column preparation" procedure available in the LC plugin

5. Open provided MS method (Bruker_diaPASEF_5min_MS.m) in timsControl and perform m/z and mobility calibration

5. Create a new sample table using the methods provided. The samples should be named as follows: date_labname_replicate (e.g. 20240207_DemoBremen_01)

6. Run 2 blanks and 10 replicate injections of 200ng K562 (injection volume: 1µl)

7. Zip each acquired data file individually and upload them
